# Various Optimization Strategies for the Isolation of Mitochondria from Sprague-Dawley Rat Liver Tissue

**DOI:** 10.1101/2025.07.30.667653

**Authors:** Chloe I. Roth, Tianyi Xia, Alyssa C. Vadovsky, Jason N. Bazil

## Abstract

Mitochondria are key organelles that establish a very large free energy drop for the ATP hydrolysis reaction in the cytoplasm of cells, which provides energy required for the cell maintenance of homeostasis. They are commonly isolated from living tissue as it makes them easier to interrogate at the biochemical level. Thus, isolated mitochondria of sufficient quality are desired to improve translational impact. To gain a greater insight into the isolation process and optimize our current isolation protocol for quality, we implemented various fine-tuning modifications to our standard homogenization, centrifugation, and purification steps with isolated rat hepatocyte mitochondria. The following modifications we tested were: i) different homogenization speeds (10,000, 14,000, and 18,000 rpm) with varying time intervals (10, 20, and 30 sec), ii) addition of an additional purification spin, and iii) use of density gradients to further purify the isolated mitochondria from non-mitochondrial contaminants. Mitochondrial quality was approximated using the well-established respiratory control ratio (RCR). The data reveal that our original protocol yields isolated mitochondria with acceptable quality and the optimization attempts produced similar or worse mitochondrial isolates. In addition, an extra purification spin decreased RCR values and therefore is not recommended. We note that the use of density gradients did not improve the RCR, but it did remove presumed peroxisomal contamination. While this protocol can be further enhanced using additional metrics, the data indicate that our current isolation protocol is sufficiently effective since additional modifications did not yield major improvements.

## INTRODUCTION

Chemical energy gradients are essential for living organisms. Growth, respiration, contraction, and even digestion, all require energy gradients generated by mitochondria. Specifically, mitochondria provide the physical, structural, and compartmental separation necessary for the biochemical reactions that create the body’s main energy source, the adenosine triphosphate (ATP) hydrolysis potential.^***1, 2***^ Although the body has various aerobic and anaerobic processes to create ATP, the most efficient way is through oxidative phosphorylation (OXPHOS) which strictly occurs in the mitochondria.^***3***^ OXPHOS is a metabolic pathway that happens in the inner mitochondrial membrane and utilizes proteins that carry, exchange, and pump different ions/molecules needed to facilitate ATP synthesis for the cell.^***4***^ OXPHOS yields 30 to 36 ATP molecules per glucose molecule, and this process is an effective energy generator and is incredibly important for maintaining life.^***5***^ As such, mitochondria are vital organelles that provide sufficient energy for metabolically active tissues such as the liver, heart, and brain.^***4***^ Without functioning mitochondria, these organs will fail, resulting in the death of the organism.^***6, 7***^

Studying mitochondria not only allows us to understand their critical role in synthesizing ATP, but also how they contribute to metabolic homeostasis^***8***^, calcium dynamics^***9, 10***^, cell differentiation^***11***^, and apoptosis.^***12***^ The process of isolating mitochondria from cells allows us to investigate these fundamental organelles in a well-controlled environment by manipulating biochemical variables involved in mitochondrial function.^***13***^ Thus, with optimized isolation protocols, we can extract mitochondria of sufficient quality as to advance the development of novel therapies for those with mitochondrial disease and disorders that lack effective treatments.^***14***^

Existing mitochondrial isolation protocols are quite similar in strategy.^***15–18***^ In general, they all harvest the desired organ, homogenize the tissue using various techniques, and remove cellular debris and undesired organelles via centrifugation. Homogenization^***19***^ is used in the mitochondrial isolation process to break the plasma membrane to release organelles into separate components that can then be further parsed by centrifugation.^***20***^ Preferences for homogenization tools differ, some using manual^***21***^ homogenizers and others electronic.^***15, 22***^ Our protocol uses an electronic homogenizer to blend and separate organelles from rat liver to test if changing the homogenization speed and time would more accurately improve mitochondrial yield during the isolation process. Centrifugation is used to separate organelles from cellular debris and undesired components in homogenized samples, which yields pure isolated mitochondria.^***23***^ The two main approaches used across mitochondrial isolation protocols are differential centrifugation (DC) and density gradient centrifugation (DG). Density gradients are typically of the sucrose^***24***^ or Percoll^***25***^ variety. DC and DG approaches present advantages and disadvantages, so both were used to investigate this for further optimization.

Current isolation protocols are more likely to utilize DC, a process that separates cellular waste and other components by size and sedimentation velocity from the mitochondria.^***26***^ DC is the fastest and most inexpensive method for isolating mitochondria; however, it is limited because it captures cellular debris that can confound mitochondrial experimental results.^***20***^ For example, peroxisome contaminants consume oxygen and can influence respiratory control ratios (RCR), or sometimes referred to as the respiratory control index.^***27***^ The RCR is a general measure of mitochondrial quality, but one caveat is that it can oversimplify data interpretation. An RCR value indicates how coupled respiration for the purpose of making ATP.^***28***^ Explicitly, it is the ratio of oxygen consumption (J_O2_) of mitochondria during OXPHOS (state 3) and LEAK (state 2 or 4).^***29***^ Peroxisomes are similar in size to mitochondria, which is why using DC can be limiting because it may not be able to effectively separate peroxisomes from the purified mitochondria. Since the RCR values are dependent on the respiration rates during different operational states, peroxisome contamination can negatively impact RCR results, decreasing its validity. This is why using the PG method could be beneficial. PG removes cellular constituents that differential centrifugation cannot.^***16***^ However, density gradients, albeit more precise, take longer and are more expensive than DC. They can also negatively impact mitochondrial function.^***30***^ Thus, to test the differences between both methods, this protocol examines the usefulness of clearing peroxisome contamination in comparison to standard DC methods.

The purpose of this study was to further modify a cardiac isolation protocol^***15, 31***^ for isolated hepatocyte mitochondria. Several variables including the homogenization speed, homogenization time, and centrifugation steps were modulated. First, the homogenization speed, homogenization duration, and whether the addition of an additional “clean up” spin resulted in improved RCR values. Next, the addition of a purification spin was implemented to improve mitochondrial quality. Then, a density centrifugation method was tested to assess whether metrics were enhanced relative to the standard differential centrifugation method. The functionality of mitochondria was then measured using the RCR as an index of comparison. In conclusion, it was found that the protocol designed for cardiac mitochondria worked efficiently without significant modification. Additionally, using the DG method removed peroxisome contaminants as expected, but did not significantly improve RCR values.

## METHODS AND PROCEDURES

Sprague Dawley (SD) rats were used according to ethical guidelines and regional laws. Animal care and handling conformed to the National Institutes of Health’s Guild for the Care and Use of Laboratory Animals and was approved by Michigan State University’s Institutional Animal Care and Use Committee. Rats (10 – 12 weeks) were euthanized after being anesthetized with 5% isoflurane and tested to be unresponsive to noxious stimuli.

Isolation Buffer (IB) consisted of 200 mM mannitol, 50 mM sucrose, 5 mM K_2_HPO_4_, 5 mM MOPS (3-(N-Morpholino) propane sulfonic acid, 4-Morpholinepropanesulfonic acid), 1 mM EGTA (ethylene glycol-bis (β-aminoethyl ether)-N, N, N′, N′-tetraacetic acid), and 0.1% (w/w) fatty-acid free BSA (bovine serum albumin). The pH was adjusted to 7.15 with 1 M KOH. Respiration Buffer (RB) consisted of 130 mM KCl, 5 mM K_2_HPO_4_, 1 mM MgCl_2_, 20 mM MOPS, and 0.1 % (w/w) fatty-acid free BSA. The pH was adjusted to 7.15 with 1 M KOH at 37 °C. A 10X IB solution was used to make Percoll™ density gradients according to the manufacturer’s instructions. Density marker beads were used (Cospheric) to form density gradients so that 1.04 g/mL was located near the middle of the 15 mL centrifuge tube. To form the continuous density gradients, the centrifuge tubes were spun at 10,000 x g for 15 min. Once formed, density gradients are stable for up to a week in the fridge. Then, on the day of isolation, 1-2 mL of the purified mitochondrial suspension was layered on top of the density gradients and spun down at 800 x g for 10 min at 4 °C. An equal volume of IB was added to the marker bead centrifugation tube to balance the centrifuge.

To extract the liver for mitochondrial purification, rats were sacrificed using a sharp guillotine after entering a deep state of isoflurane-induced anesthesia and confirmed by reflex testing. The liver was immediately removed and placed in a chilled beaker on ice. The SD rat liver contains four lobes, but three lobes yield plenty of mitochondria. The lobes were then washed with chilled IB in a 20 mL beaker to rinse off blood. Then, excess IB was removed, and the liver lobes were minced into approximately 1 mm^3^ pieces with clean scissors on ice. **Figure 1** shows a schematic of the isolation process used in this study.

**Figure 1.**
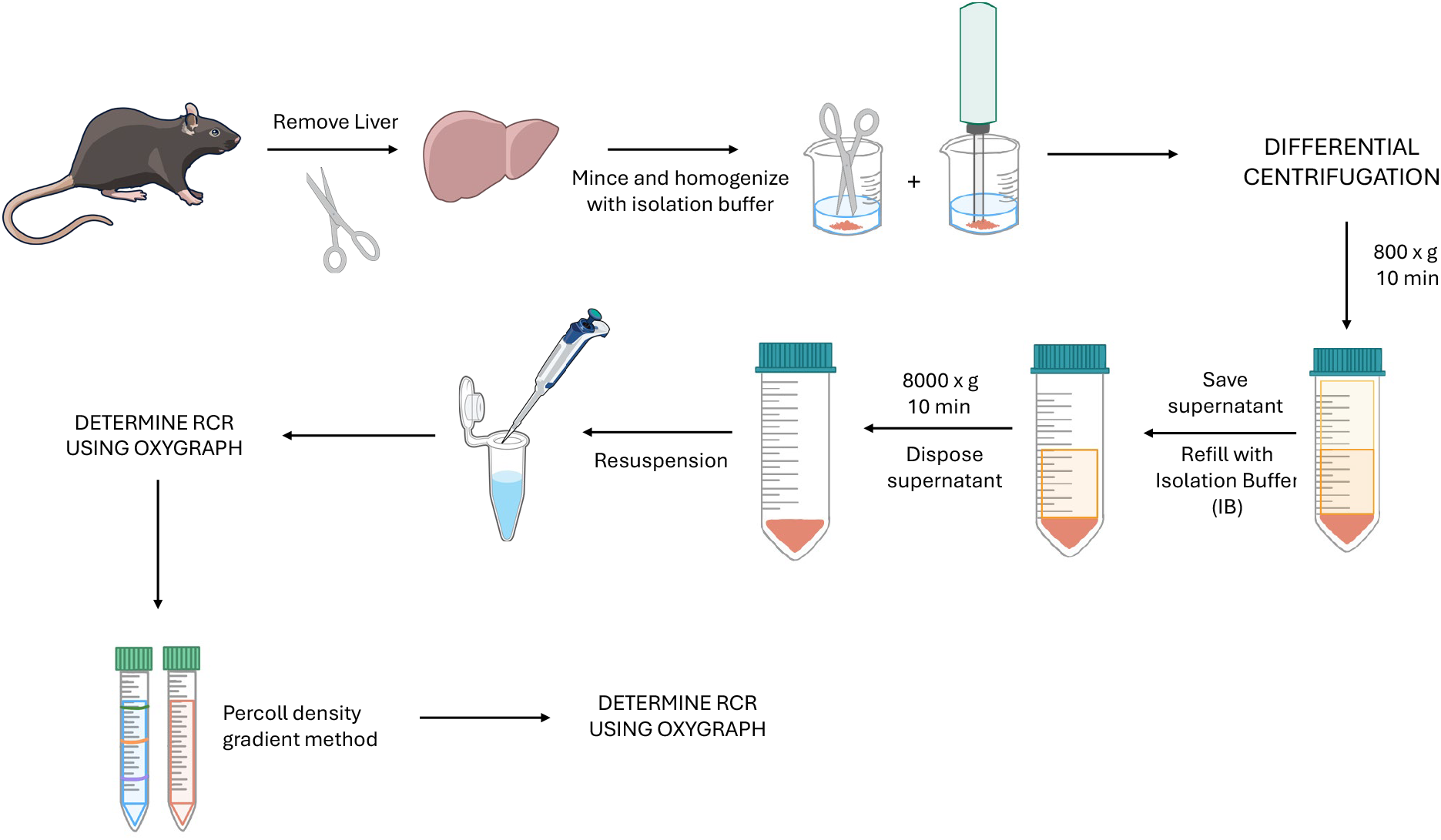
Hepatocyte Mitochondria Isolation Illustration. Step-by-step scheme of hepatocyte isolation protocol without extra clean up spin. All steps except tissue removal and respirometry were done at 4 °C. Image created through bioart.niaid.nih.gov.

The standard mitochondrial isolation protocol used in this study can be found in several previous research articles.^***15, 29, 31–33***^ In brief, tissue is minced into ~1 mm sized pieces with sharp scissors and then homogenized using an electronic homogenizer set at a speed of 18,000 rpm for 20 seconds. Following homogenization, differential centrifugation is used to purify mitochondria from the homogenate. For soft tissues, only two centrifugation steps are implemented since protease is not required for liver. The first spin is a low-speed spin at 800 x g for 10 minutes and separates cellular organelles from heavier cellular debris such as nuclei. The second spin is a high-speed spin at 8,000 x g for 10 minutes and pellets mitochondria and other similarly sized organelles such as peroxisomes. The pellet is then carefully resuspended in IB for quality control tests and experiments. Resuspension volume depends on pellet size is typically between 40 – 80 µL of IB.

For this study, the following adjustments were made to the standard isolation protocol briefly described above. First, an extra spin of 800 x g for 10 mins was done after the first 10-minute 800 x g spin. Second, the homogenization speed and time was adjusted to the following: i) 18,000 rpm for 20 sec, ii) 14,000 rpm for 26 sec, and iii) 10,000 rpm for 36 sec. These settings keep the number of shearing events approximately equal across groups. Third, the homogenization speed was set to 18,000 rpm for 10 sec, 20 sec, and 30 sec. Finally, it was tested whether continuous density gradients could improve quality control metrics for the mitochondria purified using homogenization settings of 18,000 rpm and 20 sec. **Figure 2** shows a picture and cartoon of the process and indicates how isolated mitochondria were recovered from the density gradients. Once recovered, the mitochondria were spun down at 8,000 x g for 10 min and resuspended with fresh IB.

**Figure 2.**
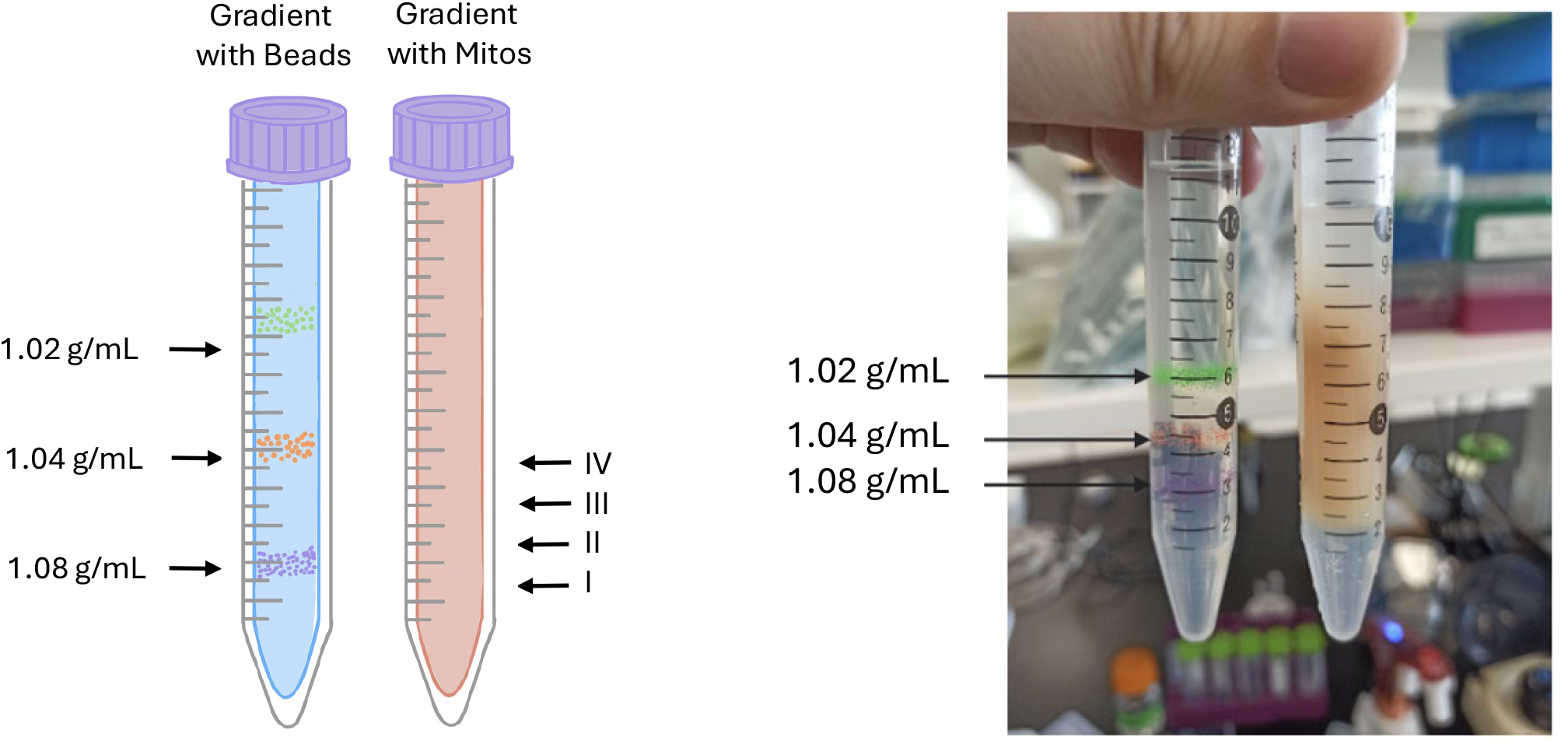
Cartoon of density gradient with marker beads and mitochondria before and after centrifugation. On the left side is a cartoon representation of the 15 mL centrifugation tubes with marker beads and isolated mitochondria. On the right side is a picture of 15 mL tubes with marker beads and mitochondrial stock after centrifugation. Roman numerals indicate the sampling locations for mitochondrial recovery. Left image created through bioart.niaid.nih.gov.

Respiration buffer was loaded into clean 2.0 mL oxygraph chambers and allowed to equilibrate with the atmosphere. Once equilibrated, the chambers were sealed and checked for bubbles. After no bubbles were detected, 10 μL of 0.2 M EGTA and 14 μL of a pyruvate/malate (1 M/0.5 M) mixture was injected into each chamber. Then 15 – 40 μL of purified mitochondria were injected and allowed to reach a steady-state for five minutes. After, 2 μL of neutralized 0.5 M ADP was injected into the chamber to initiate oxidative phosphorylation. Due to time constraints, mitochondrial protein was not quantified for this study. That said, mitochondrial respiration is linearly proportional to the number of mitochondria in the respiratory chamber, so estimates of protein load can be reasonably estimated.

### Statistical Analysis

Statistical analysis was performed in MATLAB using the anovan and ttest2 functions. The function anovan performs an n-way analysis of variance (ANOVA). The function ttest2 performs an unpaired two-sample t-test. The MATLAB documentation on the anovan function gives the following:

> “P=anovan(Y, GROUP) performs multi-way (n-way) ANOVA on the vector Y grouped by entries in the cell array GROUP. Each cell of GROUP must contain a grouping variable that can be categorical, numeric vector, character matrix, or single-column cell array of strings. Each grouping variable must have the same number of items as Y. The fitted ANOVA model includes the main effects of each grouping variable. All grouping variables are treated as fixed effects by default. The result P is a vector of p-values, one per term.”

The multcompare was used to help analyze the ANOVA output in more detail. The MATLAB documentation on this function gives the following:

> “Perform a multiple comparison of means or other estimates.
>
> multcompare performs a multiple comparison using one-way ANOVA or anocova results to determine which estimates (such as means, slopes, or intercepts) are significantly different.”

The anovan and multcompare functions were used to generate the results shown in **Figure 3**.

**Figure 3.**
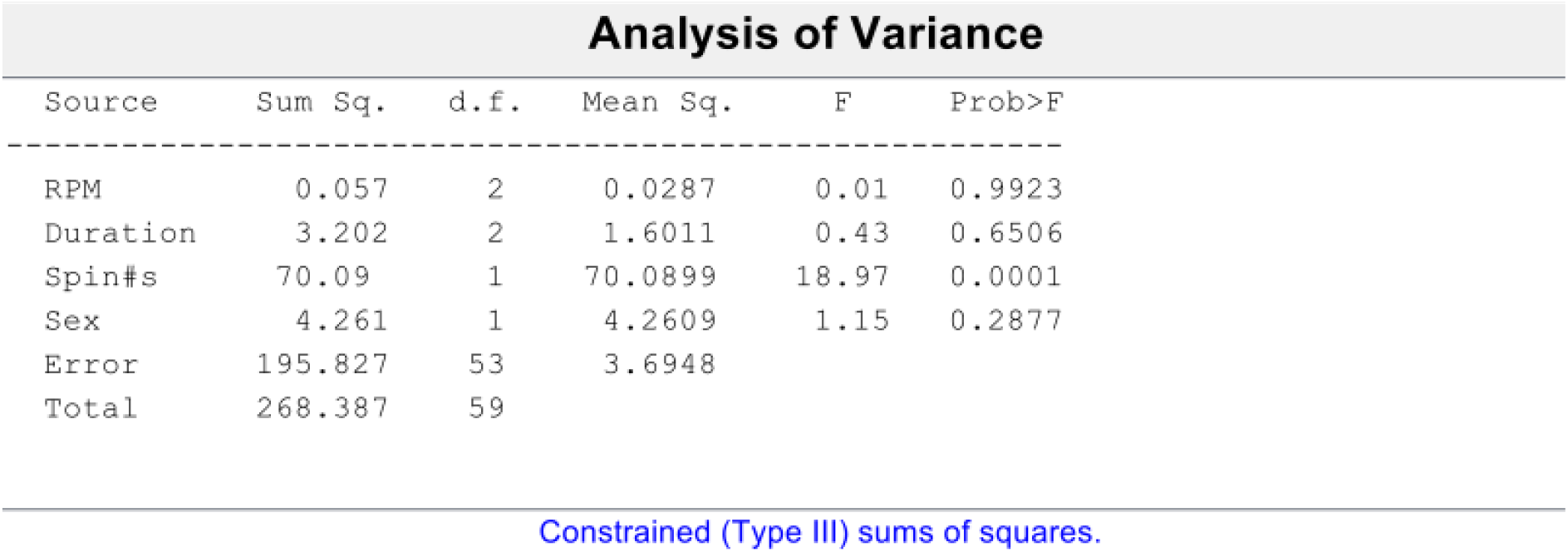
Statistical analysis results. MATLAB output for the statistical analysis performed using the anovan and multcompare functions.

From this result, the RPM had almost no effect on RCR, the Duration had a very weak effect, the Sex had a stronger but still weak effect, and the Spin#s had a strong significant effect with a p-value <0.0001. The RPM variable is for the homogenization speed (10,000, 14,000, and 18,000 rpm). The Duration variable is for the duration of homogenization (10, 20, and 30 sec). The Spin#s variable for the number of spins used to isolate mitochondria (2: slow, fast and 3: slow, slow, fast). The group-by-group results are given by the outputs from the multcompare function. Because there are 36 groups, the binomial equation gives 630 combination pairs that the multcompare function tests. Tables S1 and S2 show the multcompare results in the Supplement. The only strong statistical finding from the ANOVA was that the extra spin decreased the RCR in a consistent and reproducible manner.

## RESULTS

### Using Respiratory Control Ratio as an Index of Mitochondrial Efficiency

An RCR value indicates the quality of isolated mitochondria and how coupled respiration is to ATP production.^***7***^ Specifically, it is the ratio of mitochondrial oxygen consumption (J_O2_) during OXPHOS relative to LEAK (see **Figure 3**). Once the respiratory signal stabilizes in the presence of substrates and absence of ADP, this is referred to as the LEAK state or state 2. After ADP is added, the mitochondria begin to oxidatively phosphorylate ADP. This is called the OXPHOS state or state 3. The RCR value is calculated by dividing the OXPHOS state rate by the leak state rate. The higher the RCR value, the more coupled the mitochondria. Under healthy conditions, mitochondria in cells are well-coupled unless acted upon by signaling events, disease, or injury. Well-coupled mitochondria capture more energy from substrate oxidation and transduce it into the ATP hydrolysis potential, a potential used by nearly every reaction in the body to displace many biochemical reactions away from equilibrium and maintain homeostasis. Thus, a good mitochondrial isolation protocol will maximize the RCR value.

**Figure 4** is an illustration of the data collected from an oxygraph during the time course of a bioenergetic experiment. Substrates and EGTA were injected first. Then, mitochondria were injected into the chambers. The spike in respiration observed immediately after the addition of mitochondria is likely due to peroxisome activity.^***34***^ This was verified with data collected from the purer mitochondrial stocks after passing them through Percoll gradients (see **Figure 11**).

**Figure 4.**
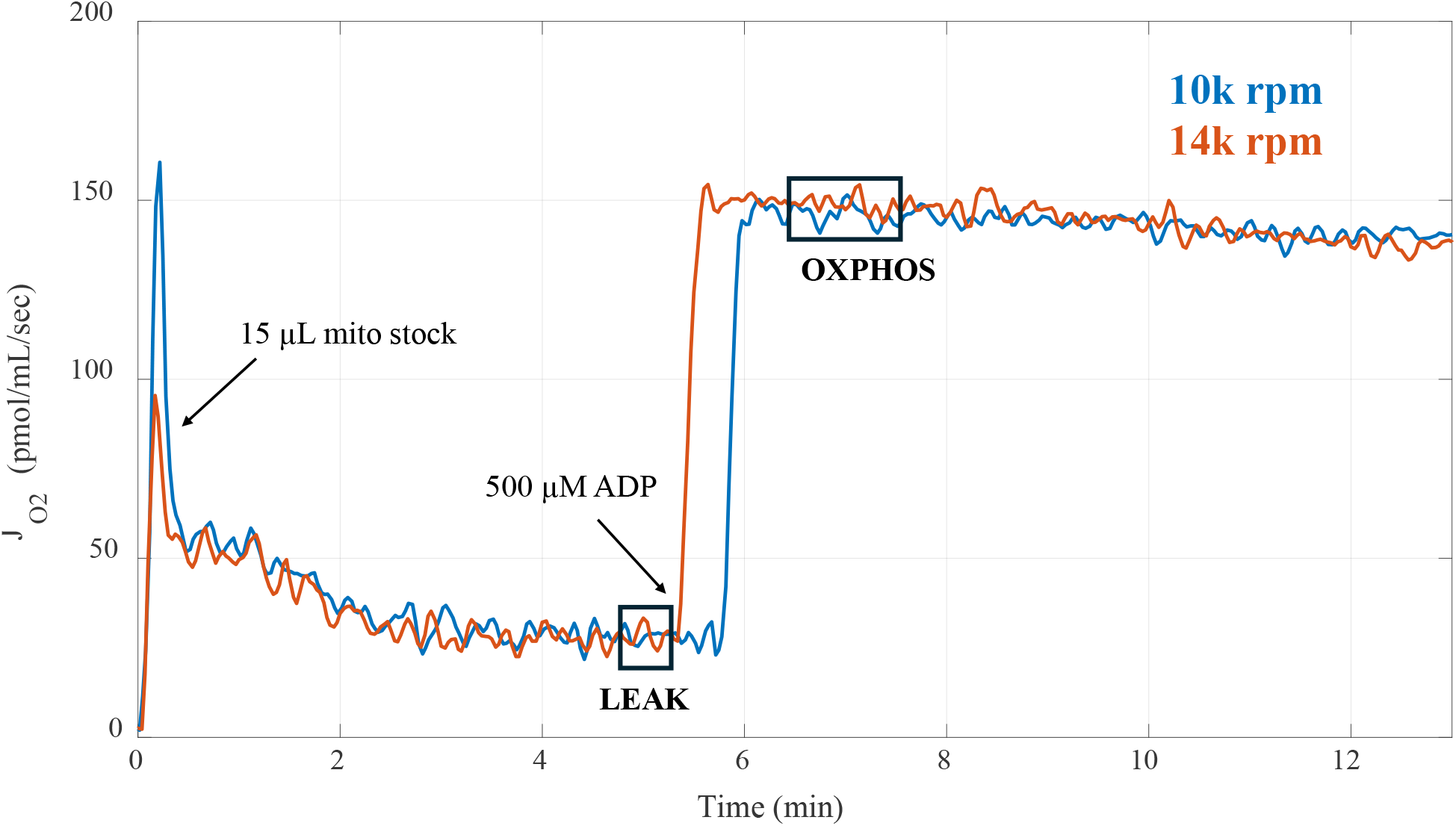
Two Chamber RCR Tracing of Isolated Hepatocyte Mitochondria using two Homogenization Speeds. Results of 15 µl of purified mitochondria injected into 2 mL of respiration buffer isolated using 10,000 rpm homogenization speed (blue) and 14,000 rpm homogenization speed (orange).

**Figure 5** displays the results of isolating mitochondria with an extra spin at different homogenizer rpm settings. The time was varied for each homogenization speed to keep shearing events relatively consistent with each sample. Based on the resulting data, there were no major differences found after changing the homogenization speed or duration. The three-spin protocol yielded RCR values worse than the standard two-spin protocol in a statistically significant manner with a p-value < 0.001.

**Figure 5.**
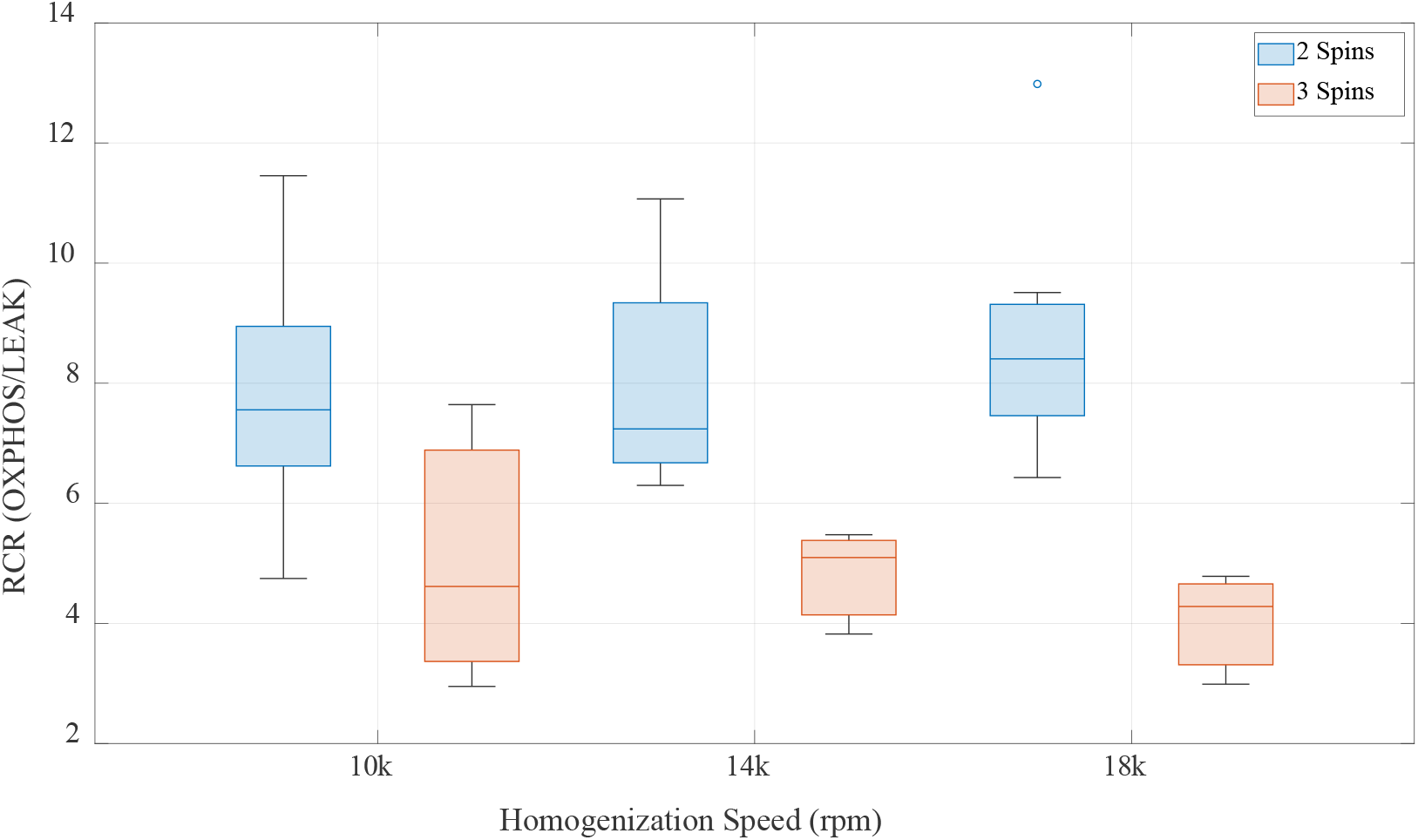
RCR Values of Isolated Hepatocyte Mitochondria with Varying Homogenization and Purification Spins. RCR value of rat liver mitochondria with 18,000 rpm for 20 seconds, 14,000 rpm for 26 seconds, and 10,000 rpm for 36 seconds without (2 spins) and with (3 spins) an additional slow spin to normal protocol. Data pooled from male and female results. Blue-edged circle in the 18k group is an outlier.

**Figure 6** shows the results of isolating mitochondria at different homogenizer rpm settings while keeping the total number of shearing events roughly equivalent. Within the same total shearing events, 18,000 rpm for 20 seconds seemed to favor female rats while 14,000 rpm for 24 seconds seemed to favor male rats. With 10,000 rpm and 36 seconds, RCR did not seem to favor any sex as there was not a significant difference between males and females. Based on the collective results, it was decided to stick with the original 18,000 rpm setting for the homogenizer during the next step shown in **Figure 8**.

**Figure 6.**
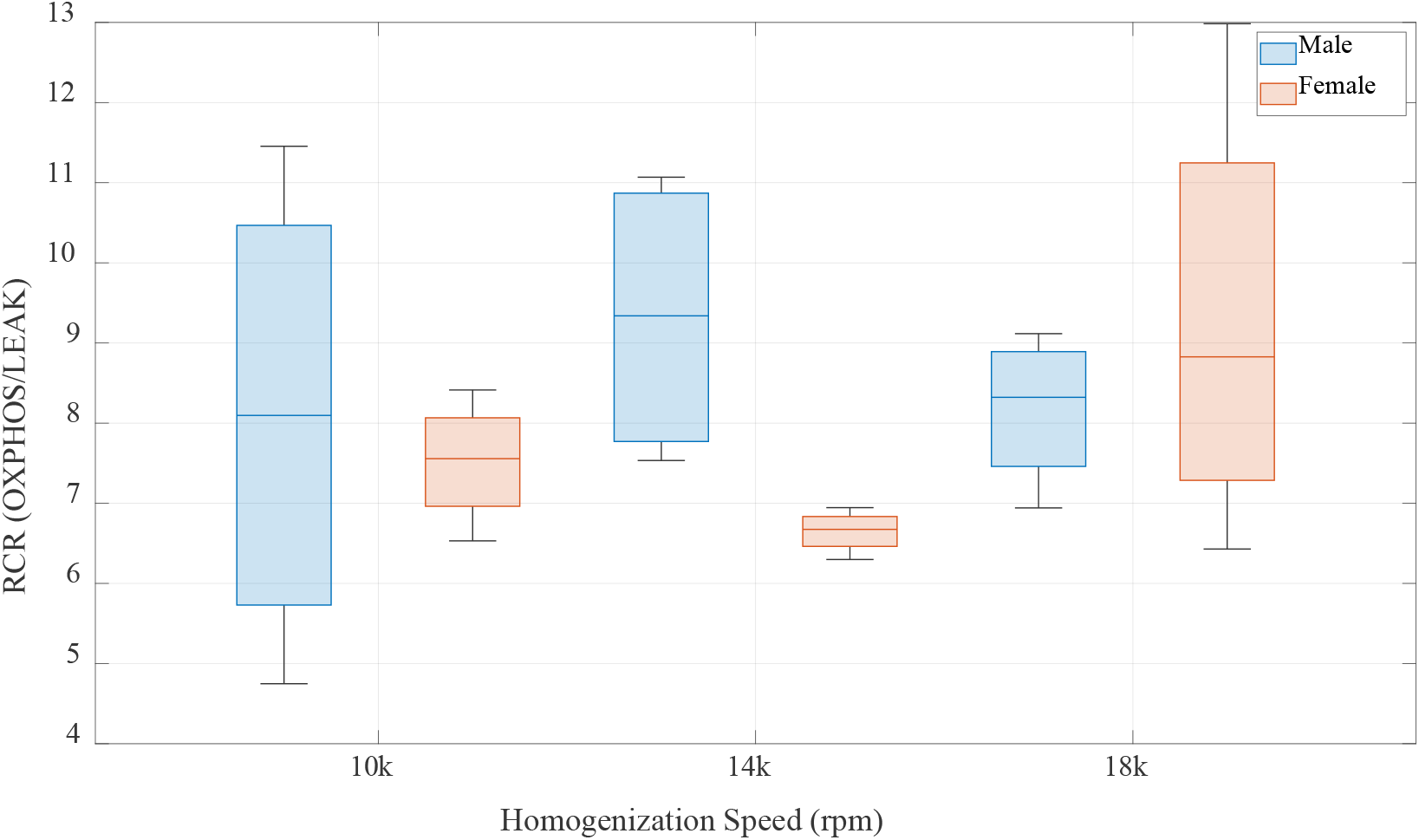
RCR Rates of Isolated Mitochondria at Three Different Homogenization Speeds (10,000 rpm, 14,000 rpm, 18,000 rpm). RCR value of both male and female rat liver mitochondria with 18,000 rpm for 20 seconds, 14,000 rpm for 26 seconds, and 10,000 rpm for 36 seconds.

**Figure 7** displays the effect of changing the number of shearing events or homogenization duration while keeping the homogenizer rpm fixed at 18,000 rpm. No strong conclusions can be drawn, so the number of shearing events does not seem to impact the quality of the mitochondrial isolates. Female rats tended to have a higher RCR than male rats for this procedure, but the results are not statistically significant. This test showed that there were no major differences between each condition that could lead to elevated RCR values.

**Figure 7.**
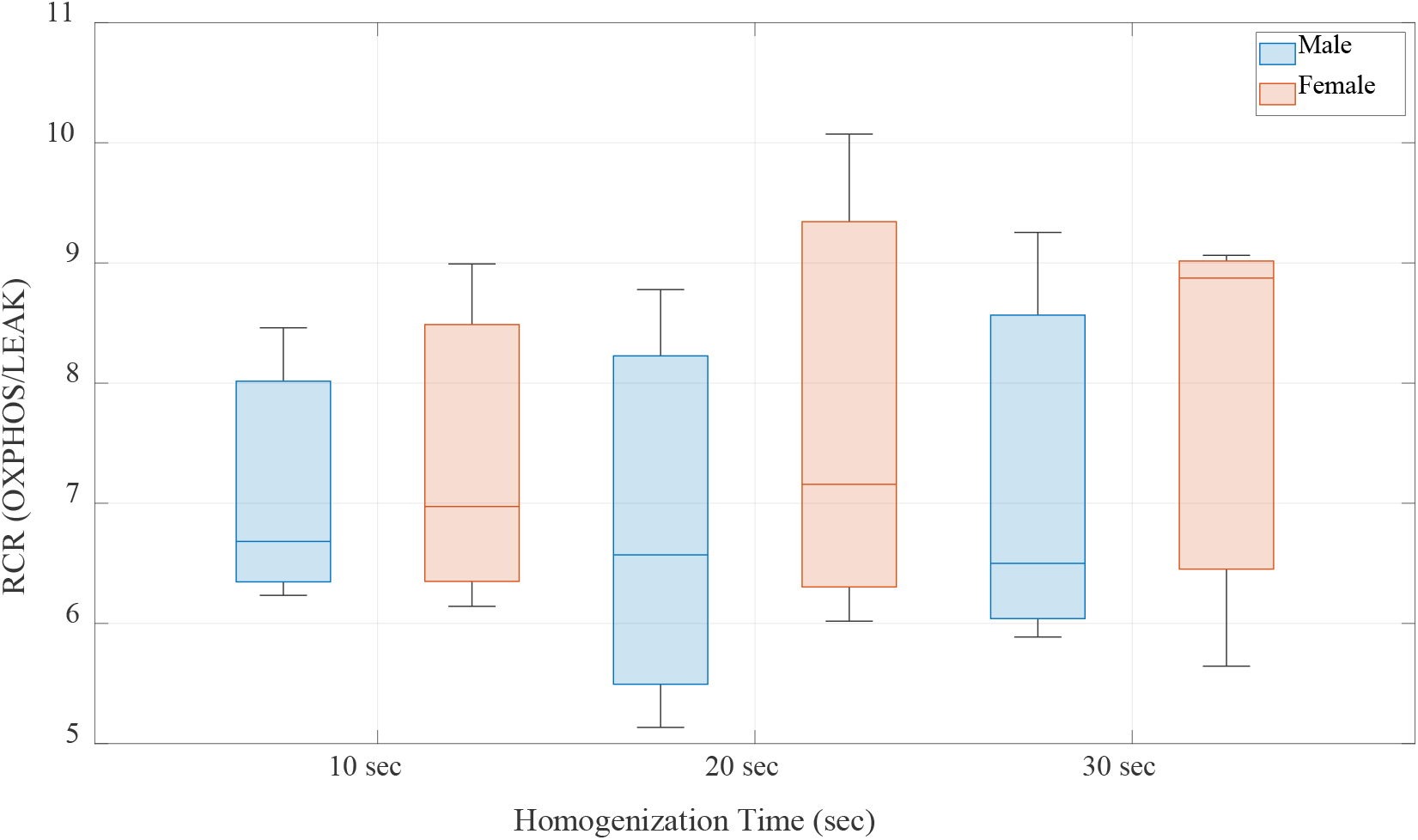
RCR values using 18,000 rpm homogenization with different time intervals. RCR results of male and female Sprague-Dawley rats of hepatocyte mitochondria with 18,000 rpm for 10 seconds, 20 seconds, and 30 seconds.

**Figure 8** shows the data of a 2:1 ratio of males and females for each condition tested. The RCR values tended to decrease after density gradient purification; however, the RCR of recovery regions II and IV (see **Figure2** for details) consisted of samples with as good or better RCR values relative to control. The recovery region that would provide the closest RCR compared to standard differential centrifugation RCR values would be between recovery regions II and IV.

**Figure 8.**
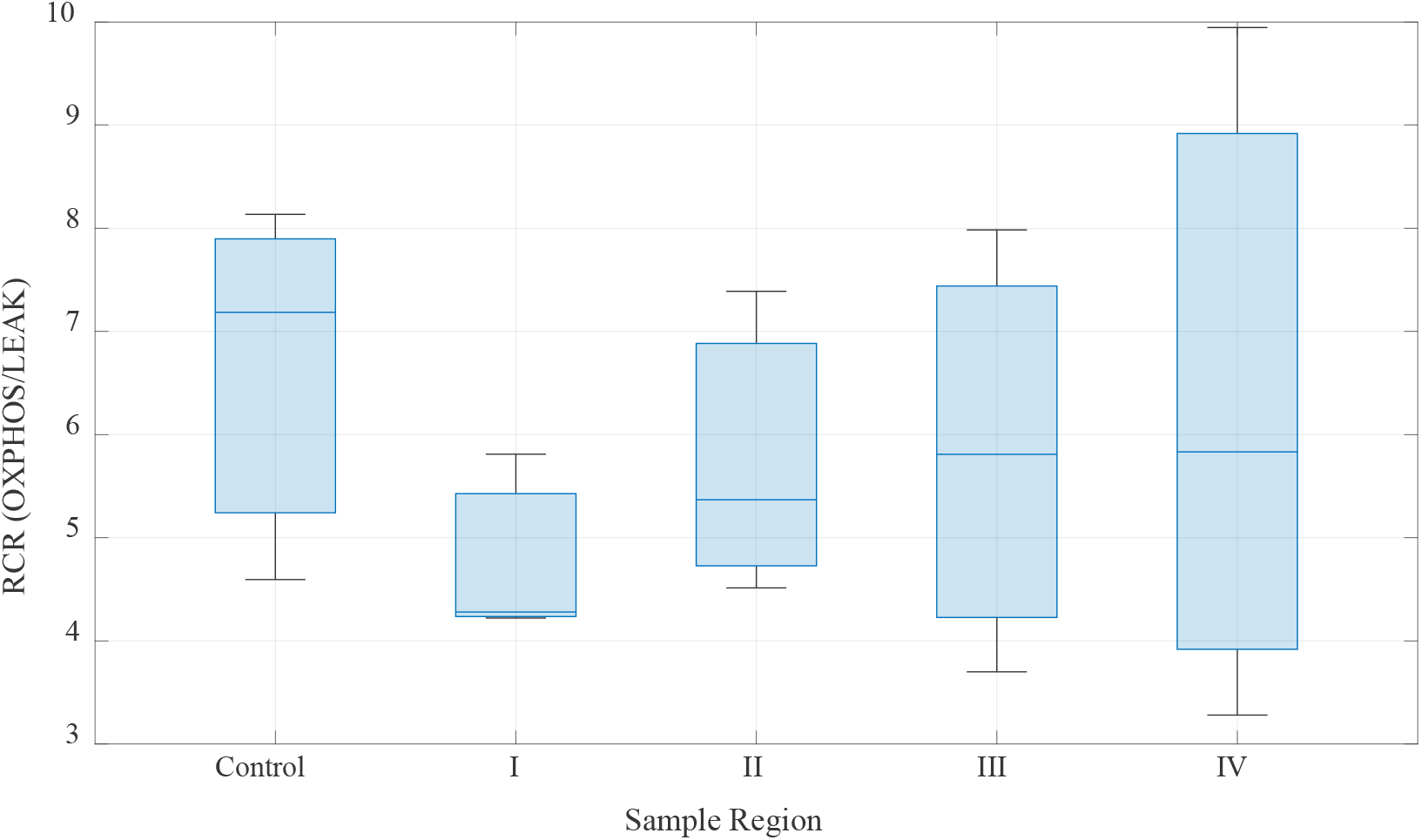
RCR of rat liver mitochondria after applying the density gradient method compared to standard differential centrifugation (Control). Quadrants define what part of the centrifugation tube isolated mitochondria were collected from. For a representative image, see **Figure2**.

Density gradients did prove to be an effective way to separate peroxisomes and mitochondria. After applying the density gradient method, the presence of an initial spike-like J_O2_ transient was absent from the bottom four regions. This was expected if peroxisomes were the source of the burst of respiratory activity when injecting the concentrated liver mitochondrial stock in the oxygraphy chambers. **Figure 9** gives an illustration of the data collected via respirometry assays during the time course of a bioenergetic experiment. The spike in respiration that was observed as before density centrifugation is significantly decreased, and in most cases entirely removed, as compared to **Figure 4**. This finding supports the beneficial uses of density gradients to remove non-mitochondrial constituents from liver mitochondrial isolates.

**Figure 9.**
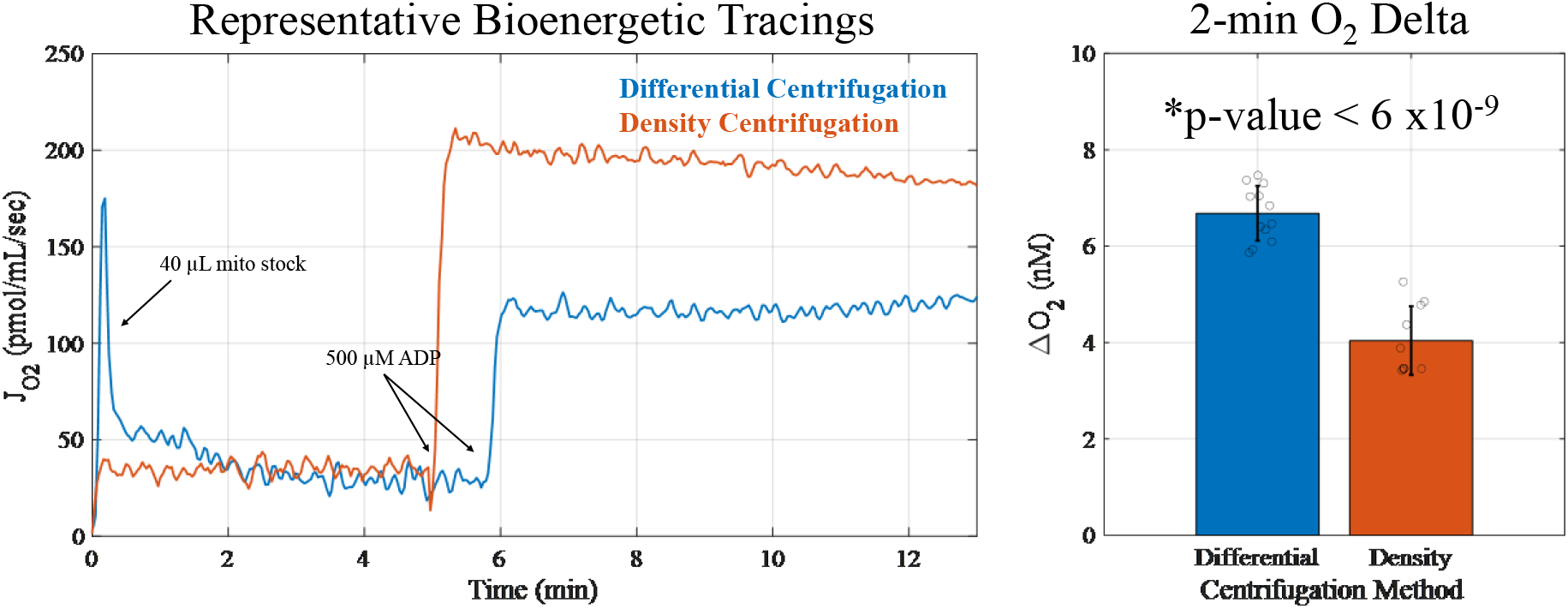
Representative bioenergetic traces showing some of the effects of density gradient centrifugation on isolated liver mitochondrial respiration. On the left, for each condition, 40 µl of either crude (blue) or density gradient purified mitochondria (orange, recovery region IV) were injected the 2 mL respiratory chambers. On the right, O_2_ consumption for each preparation was normalized to a reference LEAK state of an average across all conditions equal to 30 pmol/mL/sec and ΔO_2_ values were computed for the first two minutes of respiration after the mitochondria were injected into the respiratory chambers. This was necessary as mitochondrial protein was not determined for these sets of experiments. This is further justified due to LEAK state being directly proportional to the amount of mitochondrial protein in the chamber. LEAK was determined by time averaging the J_O2_ 30 sec before the addition of 500 µM ADP.

**Figure 10** shows a more sensitive test to determine if the PG method improved the quality of isolated mitochondria, the cyt c test. The cyt c test is a check on the integrity of the outer mitochondrial membrane. If exogenous cyt c stimulates respiratory activity during OXPHOS, the outer membrane likely becomes damaged during isolation. This damage causes the respiratory system to become rate-limited from losing cyt c. To test for this, cyt c was injected into the oxygraph chamber during state 4. This respiratory state is characterized by a high extra-mitochondrial ATP/ADP with the presence of exogenous, low activity ATPases. The percent stimulation correlates with the purity of mitochondria between the original mitochondrial stock and the different quadrants extracted from the density gradients. The expectation was less than five percent based on prior experience with cardiac mitochondria. None of the trials fell into that range and the results were generally consistent across sample region which indicates that the mitochondrial isolates did not contain subpopulations of mitochondria with variable outer membrane integrity. A more detailed analysis of the density gradient data reveals an anticipated finding that can help improve mitochondrial research efforts (see **Figure 11**).

**Figure 10.**
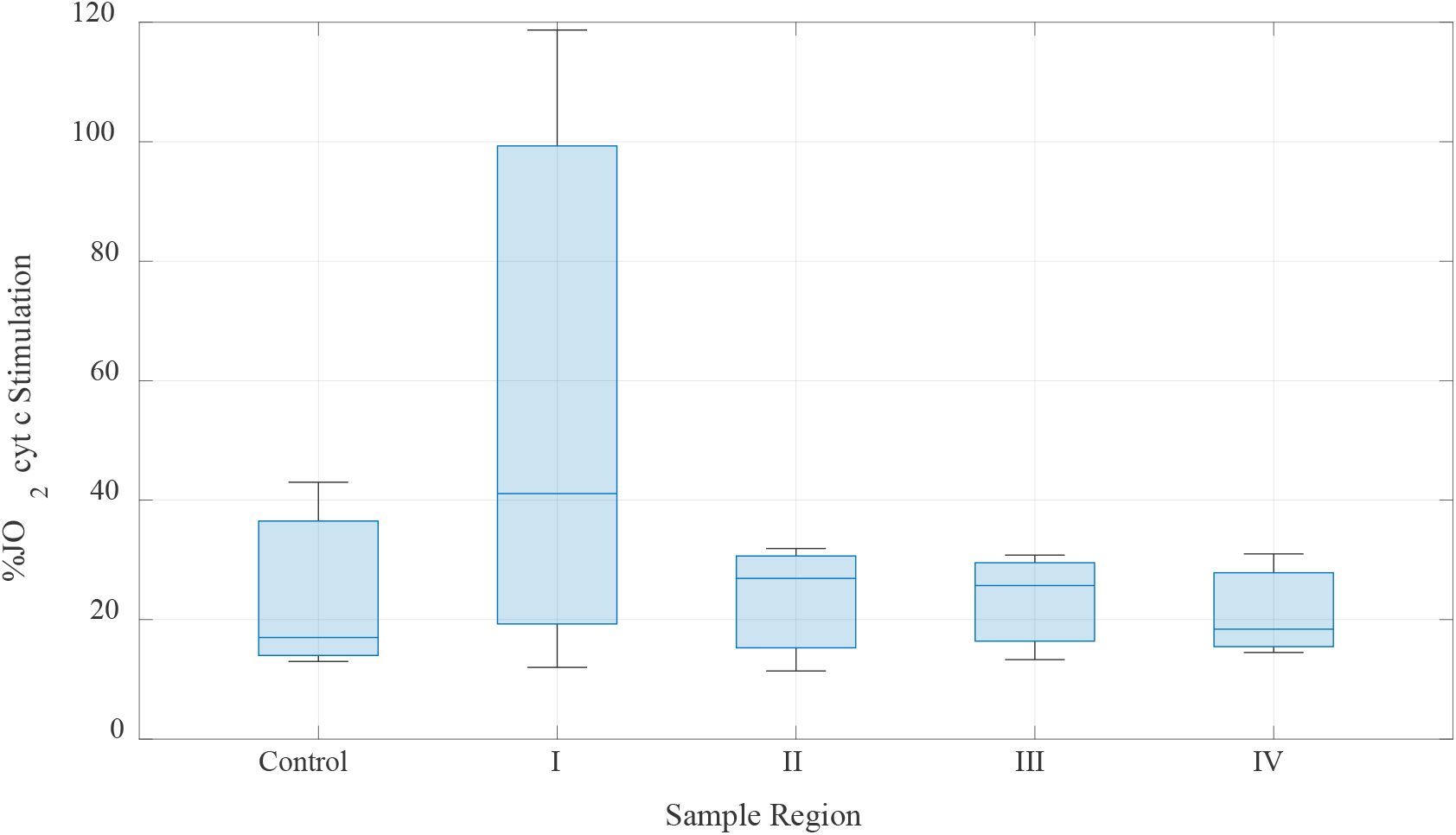
Cytochrome c test results. The effect of 10 µM cytochrome c (cyt c) on isolated mitochondria is shown as a percentage difference after applying density gradient centrifugation compared to standard differential centrifugation. Recover regions are defined as in **Figure 2**. Note, Region I had a suspected outlier which increased the upper quartile estimate.

**Figure 11.**
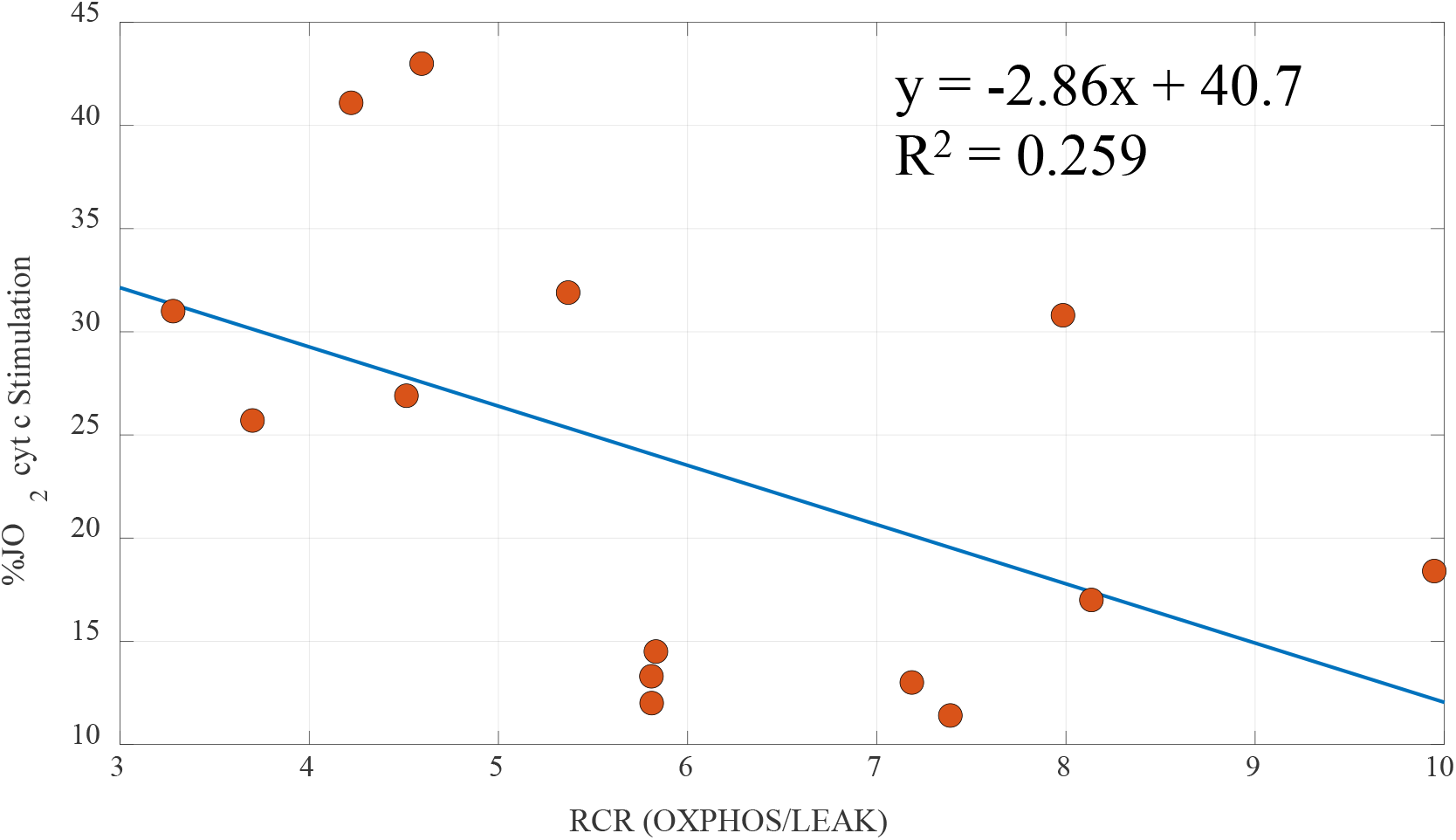
Scatter Plot of RCR value vs cytochrome c stimulation. The scatter plot displays a negative correlation between the RCR value and the cyt c stimulation rate with a linear regression line of y = −2.87x + 40.75. The data from **Figures 7** and **9** were replotted to show the relationship between RCR and the ability of cyt c to stimulate mitochondria. The suspected outlier in Figure9 was removed for this figure.

**Figure 11** shows the correlation between RCR and the percent stimulated J_O2_ after cyt c addition. The cyt c test checks the quality of the outer mitochondrial membrane, and a higher RCR generally corresponds to a lower cyt c stimulation effect. This is because a higher RCR is associated with more coupled mitochondria. Coupled respiratory control is an observation of respiratory activity and is assumed to be facilitated by a more intact outer mitochondrial membrane. Another way to see this is from a different perspective. With a damaged outer membrane, mitochondria would not be able to rapidly convert ADP into ATP unless given exogenous cyt c to fill the deficiency caused by the outer membrane damage. This is because cyt c leaks out of the IMS and cristae lumen when the outer membrane is damaged. The results indicate that mitochondria with higher RCRs tend to have smaller cytochrome c stimulation rates. While not a strong effect, a negative slope was expected and was found. These results can be used to develop more sophisticated mitochondrial quality control methods and improve experimental reproducibility.

## DISCUSSION

Based on the results from this study, a homogenization speed of 18,000 rpm with a shearing time of 20 seconds (standard protocol) resulted in RCRs than are not significantly different between the other two speeds (10,000 rpm and 14,000 rpm). These results were generated using a well-tested, standard protocol of isolating mitochondria, which further validate the procedure.^***15***^ The next step involved testing how the duration of homogenization or shearing events affected the RCR values. The results indicate that a homogenization speed of 18,000 rpm and a 20-second duration seemed to yield RCR values on par with the other durations. With gender considered as a factor that could potentially affect the RCR, sex did not have any major effect when the total shearing events were the same but seemed to play a minor but non-significant role when the shearing speeds were the same but different shearing times. More data are required before any strong inference regarding these sex-specific findings can be made.

It was expected that more pure mitochondria would be obtained by using DG since the gradient will separate damaged mitochondria and other impurities from intact mitochondria; however, the DG method had little effect on the purity of our liver mitochondrial preparations. While the RCR did not significantly improve, the location of the mitochondria in the density gradient was close to expectation coming in a bit less dense than 1.04 g/ml. The cyt c test revealed that the density gradient method produced mitochondria equal to or slightly better than mitochondria purified using standard differential centrifugation. While these results were more similar than different, there was a slightly higher RCR value as mitochondria were sampled from recovery regions II to IV. This suggests that more coupled mitochondria are more dense than less coupled ones. The density gradient method did, however, remove peroxisomes and other non-mitochondrial constituents. Thus, this method seems more apt for mitochondrial proteomic studies instead of improving the quality of isolated mitochondria.

## CONCLUSIONS

The current protocol results in high RCR values from isolated liver mitochondria. A few technical parameters were tested and found to be relatively insensitive to the quality of mitochondria as quantified by their RCR values. A homogenization speed of 18,000 rpm for 20 sec gave reasonably good RCR values; however, more replications are needed to conclusively determine this finding. A few recent papers published RCR values from small animal livers ranging between 4-6.^***14–17***^ Over 70% of our RCR values were above 6, which places this isolation protocol in the top tier rank of isolation methods. That said, improvements can still be made, so the RCR values this protocol can produce are expected to rise after further refinement. In the future, it is worth testing the use of magnetic beads with antibody coating to check if this method would yield better RCR values. Additionally, the RCR is not the only way to measure if the mitochondria are of good quality; therefore, more quality control testing should be performed. Though the sex of the rat did not seem to have a huge impact on RCR, it is still considered an important variable to test. Future studies and experiments will examine how various isolation methods impact metabolite catabolism, OXPHOS dynamics, and ROS emission profiles.

